# Allosteric Hotspots in the Main Protease of SARS-CoV-2

**DOI:** 10.1101/2020.11.06.369439

**Authors:** Léonie Strömich, Nan Wu, Mauricio Barahona, Sophia N. Yaliraki

**Affiliations:** Department of Chemistry, Imperial College London; Department of Mathematics, imperial College London

## Abstract

Inhibiting the main protease of SARS-CoV-2 is of great interest in tackling the COVID-19 pandemic caused by the virus. Most efforts have been centred on inhibiting the binding site of the enzyme. However, considering allosteric sites, distant from the active or orthosteric site, broadens the search space for drug candidates and confers the advantages of allosteric drug targeting. Here, we report the allosteric communication pathways in the main protease dimer by using two novel fully atomistic graph theoretical methods: Bond-to-bond propensity analysis, which has been previously successful in identifying allosteric sites without *a priori* knowledge in benchmark data sets, and, Markov transient analysis, which has previously aided in finding novel drug targets in catalytic protein families. We further score the highest ranking sites against random sites in similar distances through statistical bootstrapping and identify four statistically significant putative allosteric sites as good candidates for alternative drug targeting.

## 1 Introduction

The global pandemic of COVID-19 (coronavirus disease 2019) is caused by the newly identified virus SARS-CoV-2 [1, 2, 3, 4], a member of the coronavirus family of enveloped, single-stranded ribonucleic acid (RNA) viruses that also includes the virus responsible for the severe acute respiratory syndrome (SARS) epidemic of 2003 [5]. Since coronaviruses have been known to infect various animal species and share phylogenetic similarity to pathogenic human coronaviruses, the potential of health emergency events had already been noted [6]. However, their high mutation rate similarly to other RNA viruses [7] made the development of long lasting drugs challenging. Developing therapeutics against coronaviruses is of renewed interest due to the ongoing global health emergency.

One of the main approaches for targeting coronaviruses is to inhibit the enzymatic activity of their replication machinery. The main protease (M^pro^), also known as 3C-like protease (3CL^pro^), is the best characterised drug target owing to its crucial role in viral replication [8, 9, 10]. The M^pro^ is only functional as a homodimer and the central part of the active or orthosteric site is composed of a cysteine-histidine catalytic dyad [11] (see Fig. S1B) which is responsible for processing the polyproteins translated from the viral RNA [12].

The M^pro^ of the new SARS-CoV-2 shares 96% sequence similarity with that of SARS-CoV, which also extends to a high structural similarity (r.m.s deviation of 0.53 Å between *Cα* positions) [11]. Moreover, many of the residues which are important for catalytic activity, substrate binding and dimerisation are conserved between these species [13]. Nevertheless, focusing on the mutations from SARS-CoV to SARS-CoV-2, several are located at the dimer interface (for a full list see Table S1) and it has also been suggested that the mutations Thr285Ala and Ile286Leu (see Fig. S1) are responsible for a closer dimer packing [11]. Previous mutational studies on these positions in SARS-CoV M^pro^ have revealed an impact on catalytic activity [14].

Currently, the development of SARS-CoV-2 M^pro^ inhibitors [11, 15, 16, 17], similarly to designing other coronavirus M^pro^ inhibitors [18, 19, 20], focuses on blocking the orthosteric sites to disrupt viral replication (reviewed in Ullrich & Nitsche [21]). Targeting the active site enables high affinity of the drug molecules but could result in off-target-based toxicity when binding to proteins with similar active sites. Drug resistance is another major concern, especially when the active site may potentially alter owing to mutations. Targeting an allosteric site which is distal from the main binding site provides an alternative attractive solution by increasing both the range and selectivity of drugs to fine-tune protein activity without the aforementioned disadvantages. (For reviews and recent successes see Wenthur *et al*. and Cimermancic *et al*. [22, 23]). To the best of our knowledge, there is to date no indication of such putative allosteric sites of the coronavirus M^pro^s in the literature other than a recent implication of potential allosteric regulation of SARS-CoV-2 M^pro^ [24] and simulated binding events to distant areas of the protein [25]. Encouragingly, however, there have been indications of allosteric processes mediated by the extra domain in the SARS-CoV M^pro^ [26, 27, 28, 14].

Here, we focus on investigating the allosteric properties of the protease and in particular whether there are indeed any strongly connected allosteric sites to the active site that may offer alternative ways to inhibit the virus reproduction. Despite being an attractive drug alternative approach, the identification of allosteric sites remains challenging and is still often done serendipitously. Computational prediction and description of allosteric sites has become an active field of research for allosteric drug design (for reviews see [29, 30]) as it does not require the laborious and time-consuming compound screening process. For example, molecular dynamics (MD) simulations model proteins at the atomic level and the communication pathways detected can be exploited for allosteric residue and site identification [31, 32]. To alleviate the substantial computational resources required by MD simulations and the inability to explore all the required scales involved, variations of normal mode analysis (NMA) of elastic network models (ENM) are widely applied and have achieved moderate accuracy in allosteric site detection when tested on known allosteric proteins [33, 34, 35, 36]. The field of methods for allosteric pathway or site prediction is continuously growing, with new methods ranging from statistical mechanical models [37, 38] to methods based on graph theory [39]. However, even if they overcome the computational resource requirement of atomistic MD, they do so at the cost of resolution by looking at coarse-grained structure representations.

To overcome these limitations, we recently introduced a range of methods based on high resolution atomistic graph analysis which are computationally efficient while at the same time provide insights into the global effects on a protein structure without *a priori* guidance. These computational frameworks retain key physico-chemical details through the derivation of an energy-weighted atomistic protein graph from structural information which incorporates both covalent and weak interactions which are known to be important in allosteric signalling (hydrogen bonds, electrostatics and hydrophobics) through interatomic potentials [40, 41, 42]. Based on this atomistic graph, Bond-to-bond propensity analysis quantitatively shows how an energy fluctuation in a given set of bonds significantly affects any other bond in the graph and provides a measure for instantaneous connectivity. Unlike most graph or network approaches, it is formulated on the bonds or edges of the graph and thus makes a direct link between energy and flow through bonds of the system [43]. It has been shown that Bond-to-bond propensities are capable of successfully predicting allosteric sites in a wide range of proteins without any *a priori* knowledge other than the active site [43]. Of particular relevance to the homodimeric protease studied here, it has been subsequently used to show how allostery and cooperativity are intertwined in multimeric enzymes such as the well studied aspartate carbamoyltransferase (ATCase) [44]. A complementary methodology, Markov transient analysis, further sheds light on the catalytic aspects of allostery and obtains the pathways implicated in allosteric regulation through the transients of the propagation of a random walker on the node space of the atomistic graph [41]. Crucially, while most methods obtain the shortest or optimal path, the method takes into account *all* possible pathways, as allosteric communication is known to involve multiple paths [45]. In doing so, Markov transient analysis has been successful in identifying allosteric paths in caspase-1 [41] as well as previously unknown allosteric inhibitor binding sites in p90 ribosomal s6 kinase 4 (RSK4) which complemented drug repurposing in lung cancer [46]. These two methods are complementary in their application as they have been shown to provide different insights based on the underlying allosteric mechanisms: Bond-to-bond propensity analysis gives insights into the structural connectivity while Markov transient analysis is better suited for the catalytic and time scale dependent aspects of a protein.

We here showcase the application of these methodologies in the setting of COVID-19. We analysed the SARS-CoV-2 main protease and obtained Bond-to-bond propensities for all bonds as well as Markov transient half-times *t*_1/2_ for all atoms. Our results shed light on the allosteric communication patterns in the M^pro^ dimer. They further highlight the role of the interface and capture how the subtle structural changes between SARS-CoV and SARS-CoV-2 affect their dimerisation properties. By applying a rigorous scoring procedure to our results, we identify four statistically significant hotspots on the protein which are strongly connected to the active site and propose that they hold potential for allosteric regulation of the main protease. By providing guidance for allosteric drug design we hope to open a new chapter for drug targeting efforts to combat COVID-19.

## 2 Results

The first step in our graph analysis approach is the construction of an atomistic graph from a protein data bank (PDB) [47] structure. This process takes into account strong and weak interactions like hydrogen bonds, electrostatic and hydrophobic interactions (see Methods and Fig. 4). Additionally, we can incorporate water molecules, which in the case of the M^pro^ are catalytically important and known to expand the catalytic dyad to a triad [11] (see Fig. S1B). In this analysis, we use the structures of the apo form of the SARS-CoV-2 and SARS-CoV main proteases which are deposited with PDB identifier 6Y2E [11] and 2DUC [48], respectively. Once the atomistic graph is constructed, we use Bond-to-bond propensities and Markov transients to complementary explore the connectivity within the proteins when sourced from relevant residues. To achieve this, Bond-to-bond propensity explores the instantaneous strength of communication of a perturbation to every bond in the protein which allows to identify allosteric sites [43] and investigate concepts like cooperativity in multimeric proteins [44]. Markov transients exploit the time evolution of a diffusion process on the atomistic graph to identify groups of atoms which are reached the fastest (i.e. allosteric sites) or form a communication pathway [41]. By applying quantile regression we are able to quantitatively rank all bonds, atoms and subsequently residues. This allows to score the hotspots we identified and statistically prove their significance.

### 2.1 Bond-to-bond propensities validate molecular mechanism of M^pro^

Figure 1 provides detailed insights into the Bond-to-bond propensity analysis of the SARS-CoV-2 M^pro^ when sourced from the active site residues histidine 41 and cysteine 145 in both monomers. The top scoring residues (see Table S2) reveal two main areas of interest in the M^pro^. The hotspot on the back of the monomer opposite to the active site (Fig. 1A) is described in more detail in the paragraphs below. Hotspot two is located in the dimer interface and contains four residues which form salt bridges between the two monomers. Serine 1 and arginine 4 from one monomer connect to histidine 172 and glutamine 290 from the other one, respectively. Interestingly, these bonds have been found to be essential for dimer formation which in turn is required for M^pro^ activity [49, 27].

**Figure 1:**
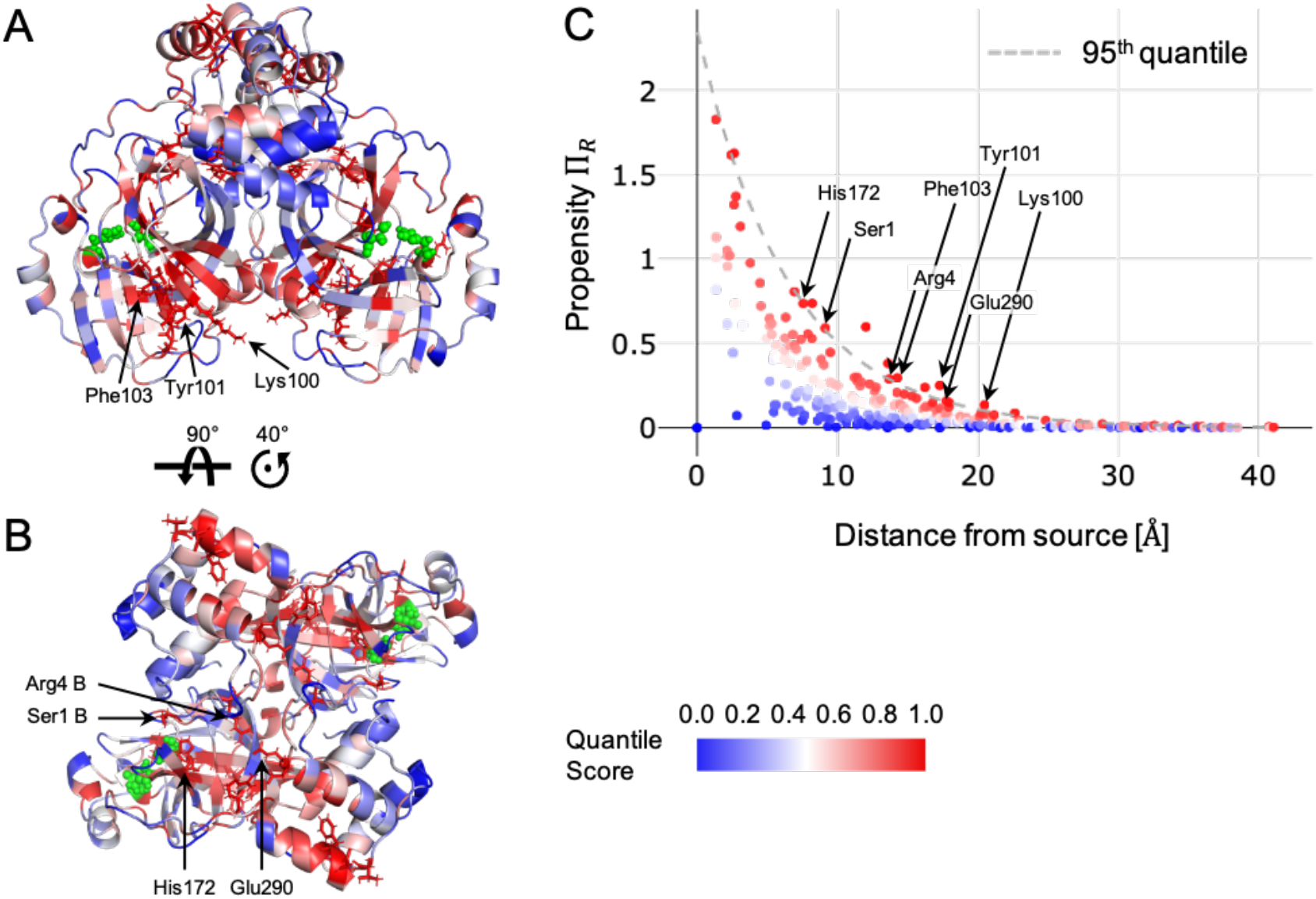
Bond-to-bond propensities of M^pro^ sourced from the orthosteric sites. The source sites have been chosen as the catalytically active residues His41 and Cys145 in both chains of the homodimer and are shown in green (front A) and top B) view). All other residues are coloured by quantile score as shown in the legend and reveal two main areas of interest with important residues labelled. C) The propensity of each residue, Π_*R*_, is plotted against the residue distance from the orthosteric site. The dashed line indicates the quantile regression estimate of the 0.95 quantile cutoff used for identifying relevant residues.

### 2.2 Protease dimerisation is under influence of mutated residues

To further clarify the interactions between the dimer halves (Fig. S1A) and how the dimer connectivity changed for the new SARS-CoV-2 protease, we ran Bond-to-bond propensity analysis sourced from two mutated residues. Alanine 285 and leucine 286 are involved in the dimer interface and have been shown to lead to a closer dimer packing when mutated from threonine 285 and isoleucine 286 in SARS-CoV [14, 11].

Hence, we chose these residues as source when looking into protease dimer connectivity in comparison between SARS-CoV-2 and SARS-CoV. Table 1 shows the top 20 residues in both structures when sorted by quantile score. We can report a strong connectivity towards dimer interface residues which is more apparent in the SARS-CoV-2 protease than in the SARS-CoV one. This can be attributed to a closer dimer packing due to the two smaller side chains of 285/286 in the new protease [11]. In a mutational study in SARS-CoV, this closer dimer packing led to an increased activity [14], however this could not be confirmed in the SARS-CoV-2 protease [11]. This was further validated when we calculated the average residue quantile score of the active site in these runs. For the active site in SARS-CoV-2 M^pro^ the score is 0.26 which is below a randomly sampled site score of 0.48 (95% CI: 0.47-0.49) and makes the active site a coldspot in this analysis. In SARS-CoV M^pro^ we detect a higher connectivity with a score of 0.50 for the active site which is nevertheless slightly above a random site score of 0.48 (95% CI: 0.47-0.48). Although we could not identify the direct link between the extra domain and the active site on an atomistic level here, we assume that studying the dimer interface residues in a systematic manner would help elucidate the link between domain III and the catalytic activity of the M^pro^.

**Table 1:**
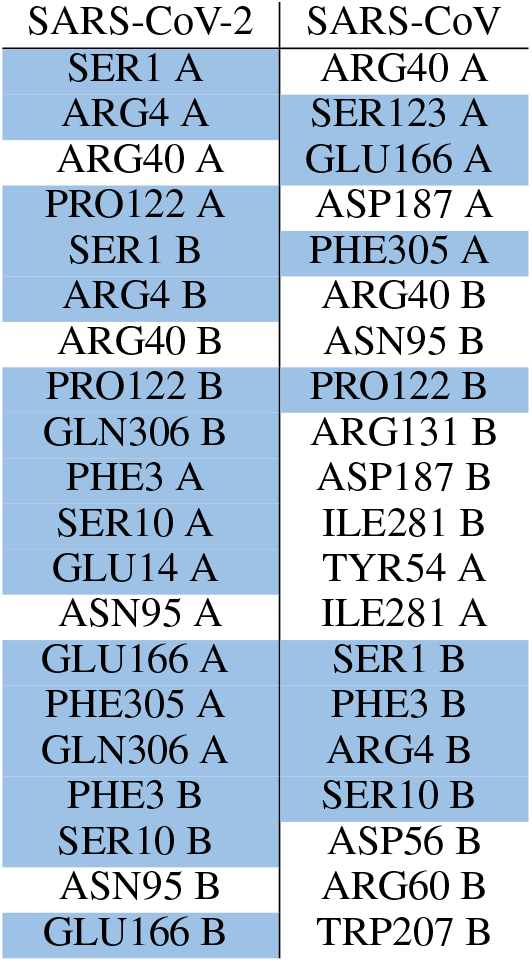
Comparison of Top 20 residues between Covid-19 and SARS main protease. Highlighted in blue are residues which are in the dimer interface.

| SARS-CoV-2 | SARS-CoV |
| --- | --- |
| SER1 A | ARG40 A |
| ARG4 A | SER123 A |
| ARG40 A | GLU166 A |
| PRO122 A | ASP187 A |
| SER1 B | PHE305 A |
| ARG4 B | ARG40 B |
| ARG40 B | ASN95 B |
| PRO122 B | PRO122 B |
| GLN306 B | ARG131 B |
| PHE3 A | ASP187 B |
| SER10 A | ILE281 B |
| GLU14 A | TYR54 A |
| ASN95 A | ILE281 A |
| GLU166 A | SER1 B |
| PHE305 A | PHE3 B |
| GLN306 A | ARG4 B |
| PHE3 B | SER10 B |
| SER10 B | ASP56 B |
| ASN95 B | ARG60 B |
| GLU166 B | TRP207 B |

### 2.3 Identification and scoring of putative allosteric sites

Bond-to-bond propensities have been shown to successfully detect allosteric sites on proteins [43] and we here present the results in the SARS-CoV-2 M^pro^ to that effect. By choosing the active site residues histidine 41 and cysteine 145 as source, we can detect areas of strong connectivity towards the active centre which allows us to reveal putative allosteric sites. We could detect two hotspots on the protease which might be targetable for allosteric regulation of the protease (Fig. 2). Most of the residues present in the two putative sites are amongst the highest scoring residues which are listed in Table S2. Site 1 (Fig. 2A shown in yellow) which is located on the back of the monomer in respect to the active site and is formed by nine residues from domain I and II (full list in Table S4). The second hotspot identified with Bond-to-bond propensities is located in the dimer interface and contains 6 residues (Tab. S5) which are located on both monomers (Fig. 2B shown in pink). Two of these residues, Glu290 and Arg4 of the respective second monomer, are forming a salt bridge which is essential for dimerisation [27]. Quantile regression allows us to rank all residues in the protein and thus we can score both sites with an average residue quantile score as listed in Table 2. Site 1 and 2 have a high score of 0.97 and 0.96, respectively and score much higher than a randomly sampled site would score with 0.53 (95% CI: 0.53-0.54) for a a site of the size of site 1 or 0.52 (95% CI: 0.51-0.53) for a site of the size of site 2.

**Figure 2:**
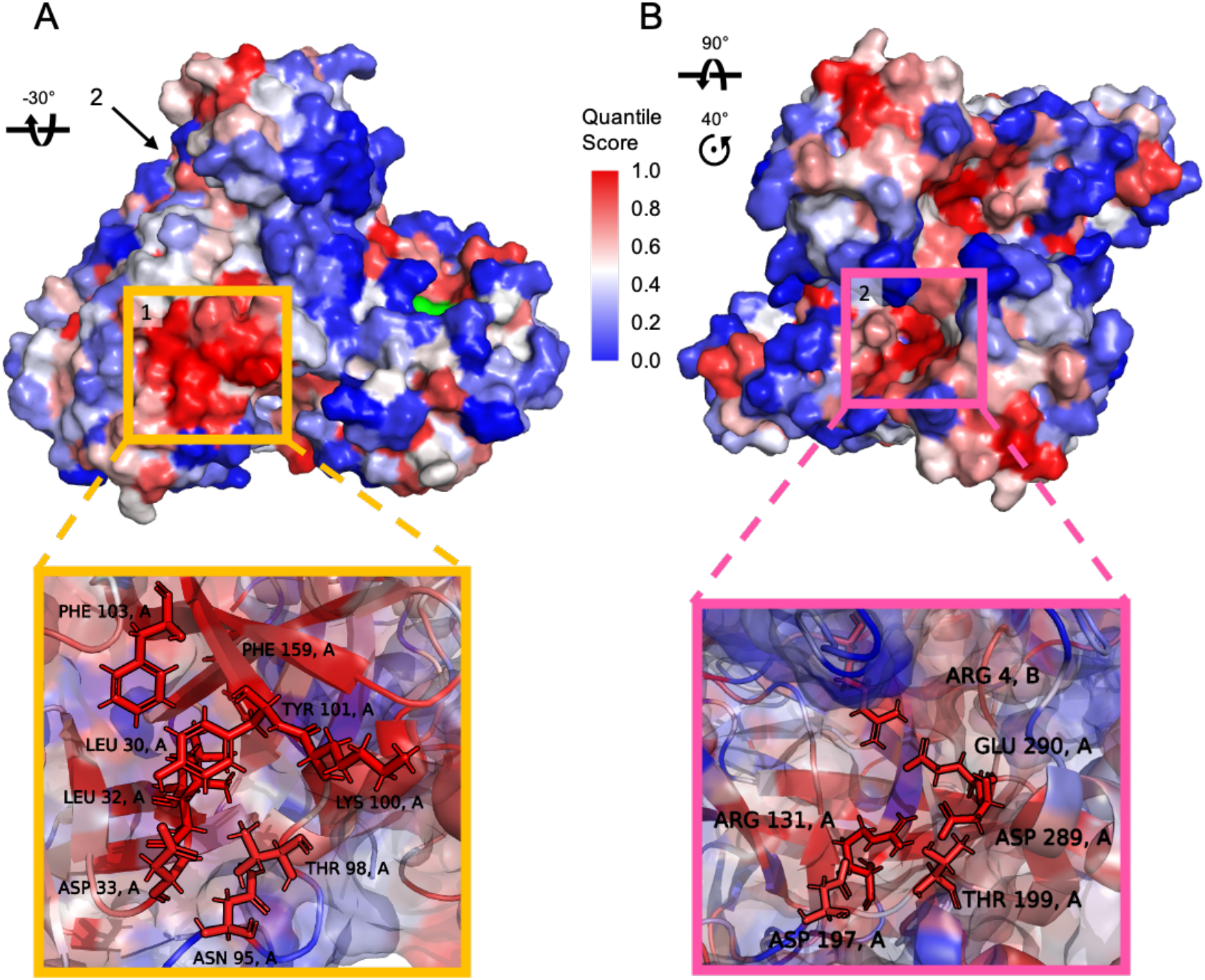
Putative allosteric sites identified by Bond-to-bond propensities. Surface representation of the M^pro^ dimer coloured by quantile score (as shown in the legend). A) Rotated front view with site 1 (yellow) which is located on the opposite of the orthosteric site (coloured in green). B) Top view with site 2 (pink) located in the dimer interface. A detailed view of both sites is provided with important residues labelled.

**Table 2:** Scoring of the 4 identified putative allosteric sites. Included is a structural bootstrap score of 1,000 randomly sampled sites with 95% confidence interval (CI).

| Site | Average Residue Quantile Score | Random Site Score [95% CI] |
| --- | --- | --- |
| Site 1 | 0.97 | 0.53 [0.53, 0.54] |
| Site 2 | 0.96 | 0.52 [0.51, 0.53] |
| Site 3 | 0.87 | 0.50 [0.49, 0.51] |
| Site 4 | 0.87 | 0.49 [0.49, 0.51] |

Our methodologies further allow to investigate the reverse analysis to assess the connectivity of the predicted allosteric sites. For this purpose, we defined the source as all residues within the respective identified sites (Tables S4 and S5). After a full Bond-to-bond propensity analysis and quantile regression to rank all residues, we are able to score the active site to obtain a measure for the connectivity towards the catalytic center (Tab. S8). For site 1 the active site score is 0.64 which is above a randomly sampled site score of 0.47 (95% CI:0.47-0.48). However, for site 2 the active site score is 0.49 which is only marginally above a randomly sampled site score of 0.48 (95% CI:0.47-0.48). As site 2 is located in the dimer interface, this is in line with the above described suggestion that the allosteric effect is not directly conferred from the dimer interface towards the catalytic centre. Nonetheless, this site might provide scope for inhibiting the M^pro^ by disrupting the dimer formation at these sites.

Overall, this missing bi directional connectivity hints to a more complex communication pattern in the protein and gave us reason to utilize another tool which has been shown to be effective in catalytic frameworks [41] like the protease. Markov transients reveal fast signal propagation which happens often along allosteric communication pathways within the protein structure. The top scoring residues with a QS > 0.95 in a Markov transient analysis sourced from the active site residues are shown in Figure 3A and a full list can be found in Table S3. In the SARS-CoV-2 M^pro^, this analysis subsequently led to the discovery of two more putative sites as shown in Figure 3C. Both hotspots are located on the back of the monomer in relation to the active site. Site 3 (shown in turquoise in Figure 3C) is located solely in domain II and consists of ten residues as listed in Table S6. One of which is a cysteine at position 156 which might provide a suitable anchor point for covalent drug design. Site 4 (orange in Figure 3C) is located further down the protein in domain I with 11 residues as listed in Table S7. Both sites were scored as described above and in the Methods section. Both sites have high average residue quantile scores of 0.87 (Tab. 2) which are significantly higher than the random site scores of 0.50 (95% CI: 0.49-0.50) and 0.49 (95% CI: 0.49-0.50), respectively.

**Figure 3:**
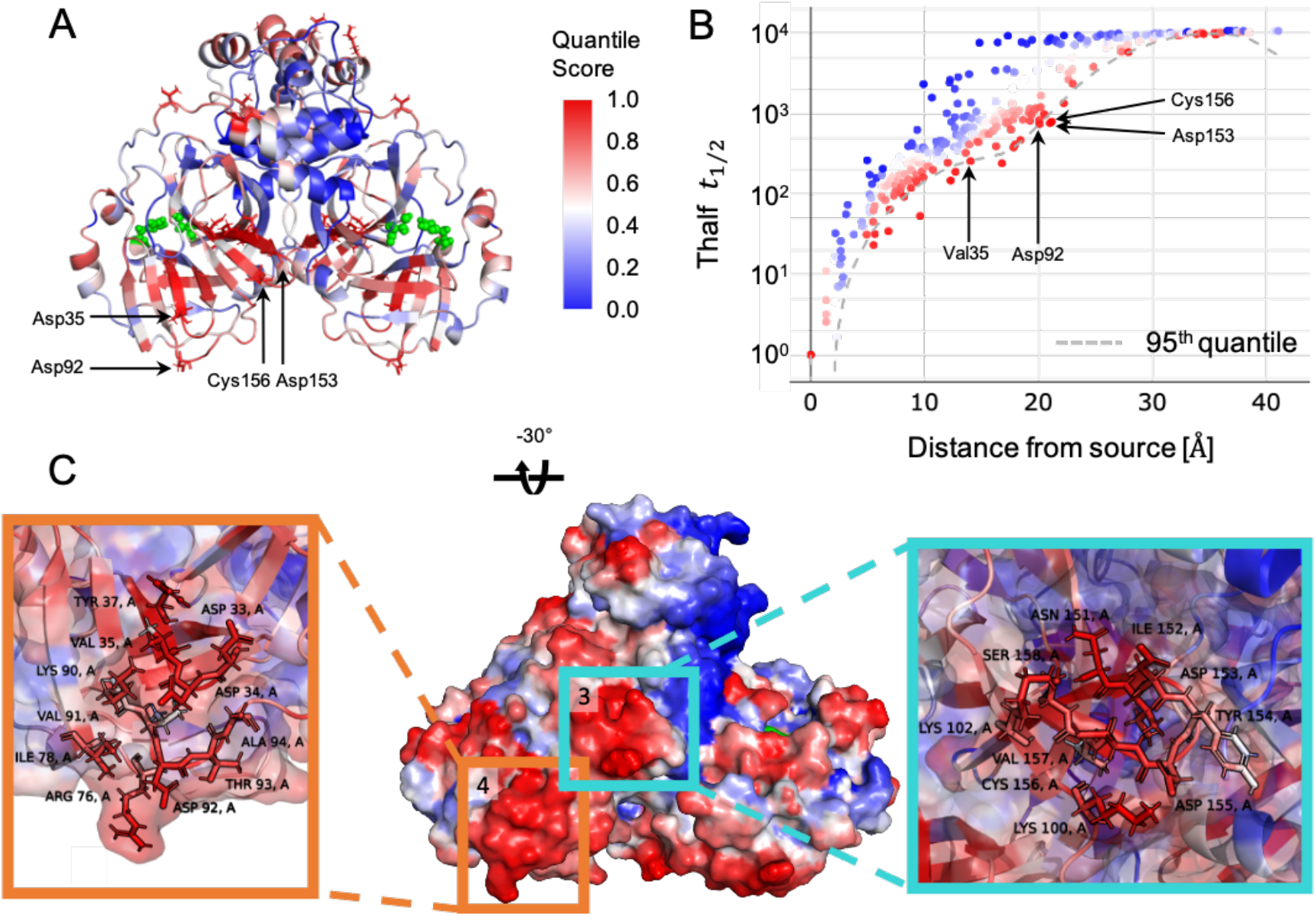
Markov transient analysis of M^pro^ sourced from the orthosteric sites. The orthosteric sites are shown in green and include His41 and Cys145 in both chains of the homodimer (front A) view). B) The *t*_1/2_ values of each residue are plotted against their distance from the orthosteric site. The dashed line indicates the quantile regression estimate of the 0.95 quantile cutoff used for identifying significant residues. The quantile scores of all residues are mapped onto the surface of the M^pro^ dimer (front A) view), coloured as shown in the legend. C) Surface representation of a rotated front view the M^pro^ dimer coloured by quantile score. Site 3 (turquoise) and 4 (orange) are located on the opposite site of the active site (coloured in green). A detailed view of both sites is provided with important residues labelled.

Following the same thought process as described for site 1 and 2, we can investigate the protein connectivity from the opposite site by sourcing our runs from the residues in site 3 and 4. We then score the active site to measure the impact of the putative sites on the catalytic centre (Tab. S8). For site 3, the active site has an average residue quantile score of 0.66 in comparison to a random site score of 0.53 (95% CI: 0.52-0.53) which indicates a significant catalytic link between site 3 and the active site. For site 4 (as for site 2) the scores are similar to a randomly sampled score, which means that we do not detect a significant connectivity from this site to the active site. Judging from previous experience in multimeric proteins this might be due to another structural or dynamic factor which we did not yet uncover between site 4 and the active site.

Overall we see a similar pattern of hot and cold spots in the SARS-CoV M^pro^ (results not shown). We find a high overlap for the identified four sites which gives us confidence, that a potential drug effort would find applications in COVID-19 as well as SARS. To provide a first indication of the druggability of the identified sites, we chose to align the fragments identified in the Diamond Light Source XChem fragment screen [50] with our sites. The screen identified 25 fragments which bind outside of the active site and 15 of these bind within 4 Å of any of the four putative allosteric sites. Due to the computational efficiency of our methodologies we were able to conduct a full analysis of all 15 structures and ran our methods from the fragments as source sites. We subsequently scored the active sites in each run (full data in Table S9) and found that the fragment deposited with the PDB identifier 5RE8 might be of particular interest as it has the highest connectivity to the active site. Moreover, one of the fragments within 4 Å of site 1 with the PDB identifier 5RGJ, has been shown to inhibit the proteolytic activity of the M^pro^ [24] and possesses a relatively high connectivity to the active site.

## 3 Discussion

During the global pandemic of COVID-19 that has started in January 2020, we have seen an increase of research activities to develop new drugs against the disease causing virus SARS-CoV-2. A wide range of approaches from chemistry, structural biology and computational modelling have been used to identify potential protease inhibitors. However, most of these initiatives focus on investigating the active site as a drug target [11, 16], high-throughput docking approaches to the active site [15] or re-purposing approved drugs [51] and protease inhibitors [52] which bind at the active site.

To increase the targetable space of the SARS-CoV-2 main protease and allow a broader approach to inhibitor discovery, we provide a full computational analysis of the protease structure which gives insights into allosteric signalling and identifies potential putative sites. Our methodologies are based on concepts from graph theory and the propagation of perturbations and fluctuations on a protein graph. We have previously demonstrated the applications of Bond-to-bond propensities and Markov transients in identifying allosteric sites and communication pathways in a range of biological settings [41, 43, 44, 46]. Applying Bond-to-bond propensities on the SARS-CoV-2 M^pro^ gave us important insights into connectivity of the protein and highlighted residues at the dimer interface. We further explored the interface residues in comparison with the SARS-CoV protease as dimerisation is known to be essential for the proteolytic activity [14] and might provide scope for inhibitor development [53]. Important for the dimer packing and mutated in SARS-CoV-2 are residues 285 and 286 [11]. When sourced from these residues, we find a higher proportion of dimer interface residues within the top 20 scoring residues for SARS-CoV-2 which confirms a stronger dimer connectivity as described in literature [11]. Although we could not identify the direct link between the mutated residues and the active site on an atomistic level here, we assume that further systematic studies of the residues at the dimer interface would provide clarity.

This gave us confidence to further explore the SARS-CoV-2 protease with our methodologies. Using the above described approaches we have identified four allosteric binding sites on the protease. We describe the location of the sites and possible implications for the proteolytic activity of the protein. Site 1 and 2 have been identified using Bond-to-bond propensities and hence have a strong instantaneous connectivity to the active site. Sourced from both sites, we noticed that site 1 is directly connected to the active site, which is detected with a score above a randomly sampled site score (0.64 > 0.47) while site 2 is indirectly connected to the active site with a active site score only slightly above that of a random site (0.49 > 0.48). This suggests that site 1 might be a functional site and any perturbation at site 1 would induce a structural change of the protease thereby impacting the active site directly. Indeed, a fragment near site 1 has been shown to exhibit some inhibitory effect on the M^pro^ in a recent study [24]. Notably, site 2, although not directly coupled to the active site as a functional site, is located in the dimer interface (Fig. 2B) and provides a deep pocket for targeting the protease and maybe disrupting dimer formation. Targeting site 2 could result in a conformation change of the protease and inhibition of dimerisation.

The sites identified with Markov transients are reached the fastest by a signal sourced from the active site and are both located at the back of each monomer in relation to the active site. Site 3 is assumed to be directly coupled to the active site as seen from the score of the active site (0.66 > 0.53) and perturbation at site 3 would thus affect the catalytic activity of M^pro^. Besides, Site 3 (Fig. 3C) contains a cysteine residue (Cys156) which provides an anchor point for covalently binding inhibitors [54]. Similar to site 2, site 4 is not directly connected to the active site. Effects exerted at site 4 could affect other parts of the protein which in turn lead to an altered activity of M^pro^.

We also include the analysis of 15 structures containing small fragments from a recent Diamond Light Source XChem fragment screen [50] which bind in proximity to the putative sites. We scored the active site (His41 and Cys145) using these fragments as the source. The active site score is analysed rigorously with a structural bootstrap to compare the effect of each fragment on the protease. Some fragments have a direct link to the active site and have been recently investigated in experimental studies [24] and might provide a first starting point for rational drug design.

Together our methods provide in depth insights into the global connectivity of the main protease. By taking our results into consideration we hope to broaden the horizon for targeting the main protease of SARS-CoV-2. This will aid in the development of effective medications for COVID-19.

## 4 Methods

### Protein Structures

We analysed the X-ray crystal structures of the apo conformations of the SARS-CoV-2 (PDB ID: 6Y2E [11]) and the SARS-CoV (PDB ID: 2DUC [48]) main proteases (M^pro^). All residues of the M^pro^ proteins that are mutated between the two viruses are listed in Table S1. Both structures contained a water molecule in proximity to the catalytic dyad formed by histidine 41 and cysteine 145. These water molecules were kept while all other solvent molecules were removed. Atom and residue, secondary structural names and numberings are in accordance with the original PDB files. The dimer interface was investigated using the online tool PDBePISA [55] (for a full list of the resulting dimer interface residues see https://doi.org/10.6084/m9.figshare.12815903).

### Atomistic Graph Construction

In contrast to most network methods for protein analysis, we derive atomistic protein graphs obtained from the three-dimensional protein structure and parameterise with physico-chemical energies, where the nodes of the graph are the atoms and the weighted edges represent interactions, both covalent bonds and weak interactions, including hydrophobic, hydrogen bonds and salt bridges (See Fig. 4). Details of this approach can be found in Refs [40, 41, 43]. We summarise the main features below and note three additional improvements, namely, in the stand-alone detection of edges without need of third-party software, the many-body detection of hydrophobic edges across scales, and, the computational efficiency of the code. For further details for the atomistic graph construction used in this work see [56, 42].

**Figure 4:**
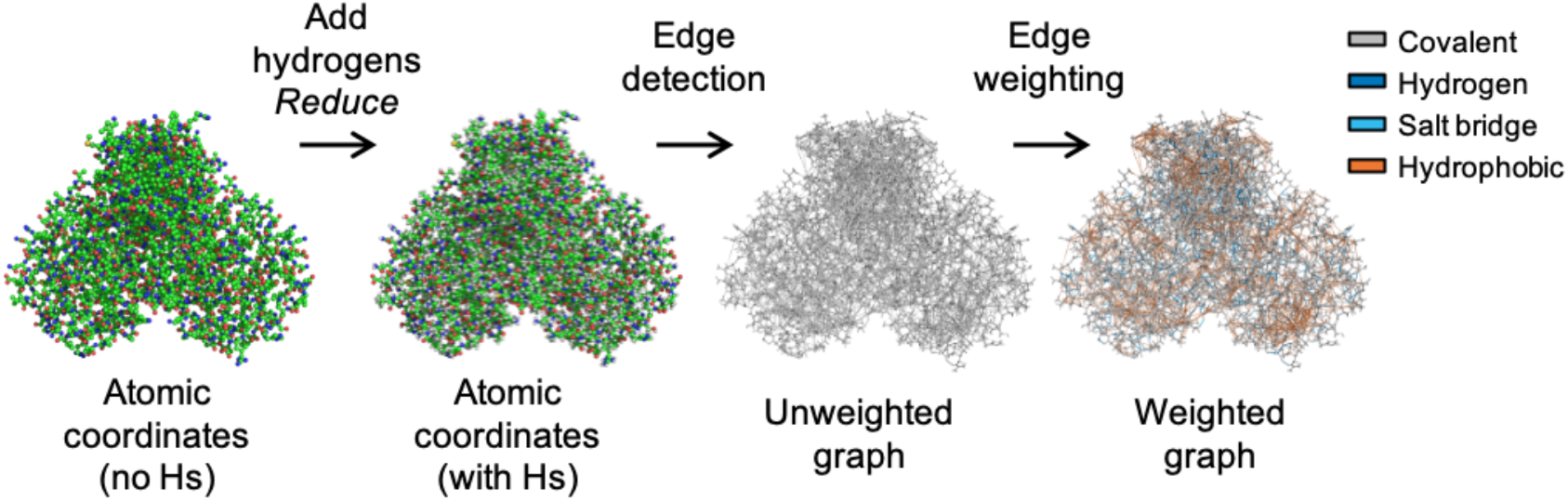
Atomistic Graph Construction. We showcase the general procedure here on the main protease of SARS-Cov-2: Atomic coordinates are obtained from the PDB (ID: 6Y2E [11] and hydrogens are added by Reduce [57]. Edges are identified and the weights are assigned, as described in the methods section, by taking into account covalent bonds as well as weak interactions: hydrogen bonds, electrostatic interactions and the hydrophobic effect which are coloured as indicated.

Figure 4 gives an overview of the workflow where we start from atomistic cartesian coordinates from PDB files. Since X-ray structures do not include hydrogen atoms and NMR structures may not report all of them, we use *Reduce* [57] to add any missing hydrogens. Hydrophobic interactions and hydrogen bonds are identified with a cutoff of 8 Å and 0.01 kcal/mol respectively. The edges are weighted by their energies: covalent bond energies from their bond-dissociation energies [58], hydrogen bonds and salt bridges by the modified Mayo potential [59, 60] and hydrophobic interactions are calculated using a hydrophobic potential of mean force [61].

### Bond-to-bond Propensities

Bond-to-bond propensity analysis was first introduced in Ref. [43] and further discussed in Ref. [44], hence we only briefly summarise it here. This edge-space measure examines and exhibits the instantaneous communication of a perturbation at a source towards every bond in the protein. The edge-to-edge transfer matrix *M* was introduced to study non-local edge-coupling and flow redistribution in graphs [62] and an alternative interpretation of M as a Green function is employed to analyse the atomistic protein graph. The element *M_ij_* describes the effect that a perturbation at edge *i* has on edge *j*. *M* is given by

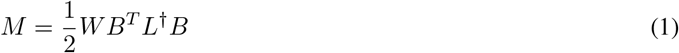

where B is the *n* x *m* incidence matrix for the atomistic protein graph with *n* nodes and *m* edges; *W* = diag(*w_ij_*) is an *m* x *m* diagonal matrix which possesses all the edge interaction energies with *w_ij_* as the weight of the edge connecting nodes i and j, i.e. the bond energy between the atoms; and L^†^ is the pseudo-inverse of the weighted graph Laplacian matrix *L* [63] and defines the diffusion dynamics on the energy-weighted graph [64].

To evaluate the effect of perturbations from a group of bonds *b′* (i.e., the source), on bond *b* of other parts of the protein, we define the bond propensity as:

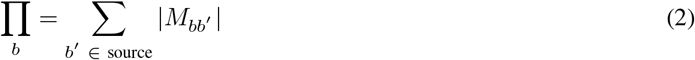

and then calculate the residue propensity of a residue *R*:

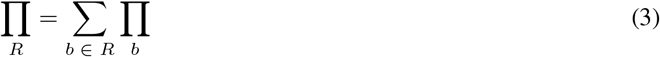

### Markov Transient Analysis (MTA)

A complementary, node-based method, Markov Transient analysis (MTA) identifies areas of the protein that are significantly connected to a site of interest, the source, such as the active site, and obtains the signal propagation that connects the two sites at the atomistic level. The method has been introduced and discussed in detail in Ref. [41] and has successfully identified allosteric hotspots and pathways without any *a priori* knowledge [41, 46]. Importantly, it captures *all* paths that connect the two sites. The contribution of each atom in the communication pathway between the active site and all other sites in a protein or protein complex is measured by the characteristic transient time *t*_1/2_,

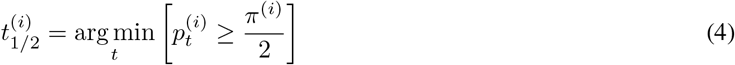

where 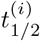 is the number of time steps in which the probability of a random walker to be at node *i* reaches half the stationary distribution value. This provides a measure of the speed by which perturbations originating from the active site diffuse into the rest of the protein by a random walk on the above described atomistic protein graph. To obtain the transient time *t*_1/2_ for each residue, we take the average *t*_1/2_ over all atoms of the respective residue.

### Quantile Regression (QR)

To determine the significant bonds with high bond-to-bond propensity and atoms with fast transient times *t*_1/2_ at the same geometric distance from the source, we use conditional quantile regression (QR) [65], a robust statistical measure widely used in different areas [66]. In contrast to standard least squares regressions, QR provides models for conditional quantile functions. This is significant here because it allows us to identify not the “average” atom or bond but those that are outliers from all those found at the same distance from the active site and because we are looking at the tails of highly non-normal distributions.

As the distribution of propensities over distance follows an exponential decay, we use a linear function of the logarithm of propensities when performing QR while in the case of transient times which do not follow a particular parametric dependence on distance, we use cubic splines to retain flexibility. From the estimated quantile regression functions, we can then compute the quantile score for each atom or bond. To obtain residue quantile scores, we use the minimum distance between each atom of a residue and those of the source. Further details of this approach for Bond-to-bond propensities can be found in Ref. [43] and for Markov Transient Analysis in Ref. [67].

### Site scoring with structural bootstrap sampling

To allow an assessment of the statistical significance of a site of interest, we score the site against 1000 randomly sampled sites of the same size. For this purpose, the average residue quantile score of the site of interest is calculated. After sampling 1000 random sites on the protein, the average residue quantile scores are calculated. By performing a bootstrap with 10,000 resamples with replacement on the random sites average residue quantile scores, we are able to provide a confidence interval to assess the statistical significance of the site of interest score in relation to the random site score.

### Residues used when scoring the active site

For scoring the active site as a measure of the connectivity towards the main binding site, we use all non-covalent hits bound in the active site from the XChem fragment screen against the SARS-CoV-2 M^pro^ [50]. The 22 found structures were further investigated using PyMol v.2.3 [68] for residues which have atoms within 4Å of any of the bound fragments. These residues are Thr25, Thr26, His41, Cys44, Thr45, Ser46, Met49, Tyr54, Phe140, Leu141, Asn142, Ser144, Cys145, Met162, His163, His164, Met165, Glu166, Leu167, Pro168, Asp187, Arg188, Gln189, Thr190 and constitute the active site as a site of interest in all scoring calculations.

### XChem fragment screen hits selection

From the above mentioned XChem fragment screen against the SARS-CoV-2 M^pro^ [50], 25 hits were found at regions other than the active site. The 15 fragments which contain atoms that are within 4Å from any of the putative allosteric site residues we obtained were selected as candidates for further investigation as shown in Table 3.

**Table 3:** XChem fragments in 4 Å proximity to the identified allosteric sites.

| Site | Fragment PDB ID |
| --- | --- |
| Site 1 | 5RGJ, 5RE8, 5RF4, 5RF9, 5RFD, 5RED, 5REI, 5RF5, 5RGR |
| Site 2 | 5RF0, 5RGQ |
| Site 3 | 5RF9 |
| Site 4 | 5RGG, 5RE5, 5RE7, 5RFC, 5RE8, 5RF4, 5RFD |

For each of these fragment-bound structures, we performed Bond-to-bond propensity and Markov transient analyses to evaluate the connectivity to the active site. The active site was scored as described above.

### Visualisation and Solvent Accessible Surface Area

We use PyMol (v.2.3) [68] for structure visualisation and presentation of Markov transient and Bond-to-bond propensity results directly on the structure. The tool was also used to calculate the residue solvent accessible surface area (SASA) reported here, with a rolling probe radius of 1.4 and a sampling density of 2.

## Data availability

All data presented in this study are available at figshare with DOI: 10.6084/m9.figshare.12815903.

## Acknowledgements

We acknowledge helpful discussions with Florian Song, Francesca Vianello, Ching Ching Lam and Jerzy Pilipczuk. This work was funded by a Wellcome Trust studentship to L.S. [grant number 215360/Z/19/Z]. M.B. and S.N.Y. acknowledge funding from the EPSRC award EP/N014529/1 supporting the EPSRC Centre for Mathematics of Precision Healthcare.

## Author contributions

L.S., N.W., M.B and S.N.Y. conceived the study. L.S and N.W. performed the computations, L.S. created the figures and all authors analysed the data and wrote the manuscript.

