## Supplemental Information for "Allosteric Hotspots in the Main Protease of SARS-CoV-2"

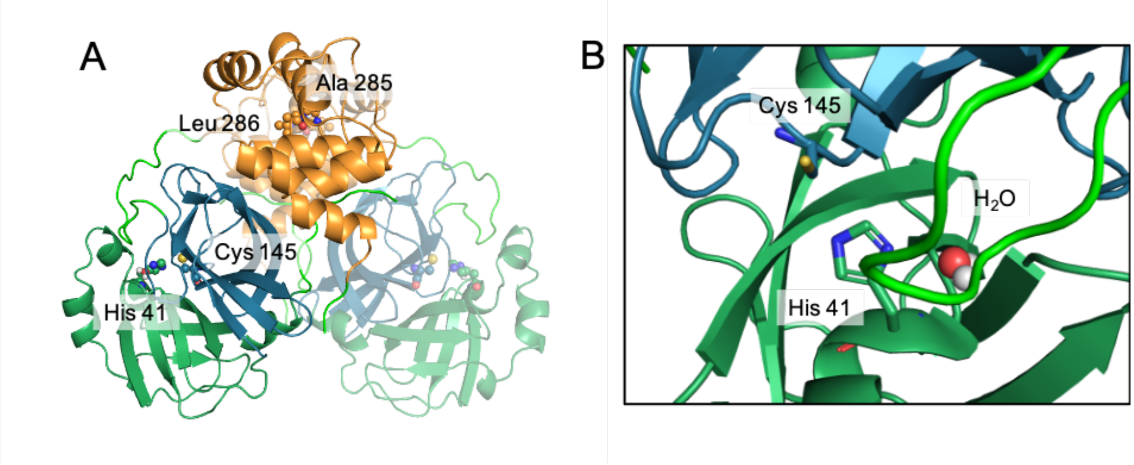


**Figure S1: Overview of the SARS-CoV-2 main protease dimer.** Atomic coordinates are obtained from the PDB (ID: 6Y2E). A) shows the full dimer with the active site residues on both monomers shown as spheres. The second monomer is shown in less colour to visualise where the monomers interact. Colours are according to domain: Domain I residues 10 to 99 - dark green, domain II residues 100 to 182 - dark blue, domain III residues 198 to 303 - orange, loops in light green. B) Zoom in of the active site with histidine 41 and cysteine 145 forming a catalytic dyad which is extended by a water molecule in close proximity.

**Table S1: Mutated residues from SARS-CoV to SARS-CoV-2.**

| SARS-CoV | Residue Number | SARS-CoV-2 |
|---|---|---|
| THR | 35 | VAL |
| ALA | 46 | SER |
| SER | 65 | ASN |
| LEU | 86 | VAL |
| ARG | 88 | LYS |
| SER | 94 | ALA |
| HIS | 134 | PHE |
| LYS | 180 | ASN |
| LEU | 202 | VAL |
| ALA | 267 | SER |
| THR | 285 | ALA |
| ILE | 286 | LEU |

**Table S2: Top scoring residues of Bond-to-bond propensity analysis in SARS-CoV-2 $M^{pro}$.** All residues are ranked by quantile score (QS $\geq 0.95$) and are shown with residue name, residue number and chain ID.

| Residue | Quantile Score |
|---|---|
| LEU32 A | 1 |
| LYS100 A | 1 |
| TYR101 A | 1 |
| ARG131 A | 1 |
| PHE159 A | 1 |
| LEU32 B | 1 |
| LYS100 B | 1 |
| TYR101 B | 1 |
| PHE159 B | 1 |
| LEU30 A | 0.98 |
| PHE103 A | 0.98 |
| HIS163 A | 0.98 |
| ASP229 A | 0.98 |
| ASP289 A | 0.98 |
| LEU30 B | 0.98 |
| PHE103 B | 0.98 |
| ARG131 B | 0.98 |
| HIS163 B | 0.98 |
| ASP289 B | 0.98 |
| SER1 A | 0.97 |
| ARG4 A | 0.97 |
| LEU27 A | 0.97 |
| HIS172 A | 0.97 |
| PHE230 A | 0.97 |
| GLU290 A | 0.97 |
| SER1 B | 0.97 |
| ARG4 B | 0.97 |
| ASP33 B | 0.97 |
| HIS172 B | 0.97 |
| ASP229 B | 0.97 |
| PHE230 B | 0.97 |
| GLU290 B | 0.97 |
| VAL20 A | 0.95 |
| ASP33 A | 0.95 |
| THR226 A | 0.95 |
| LEU272 A | 0.95 |
| VAL20 B | 0.95 |
| LEU27 B | 0.95 |
| ARG40 B | 0.95 |
| THR226 B | 0.95 |

**Table S3: Top scoring residues of Markov transients analysis in SARS-CoV-2 $M^{pro}$.** All residues are ranked by quantile score (QS $\geq 0.95$) and are shown with residue name, residue number and chain ID.

| Residue | Quantile Score |
|---|---|
| PHE150 A | 1 |
| CYS156 A | 1 |
| THR201 A | 1 |
| PHE150 B | 1 |
| CYS156 B | 1 |
| GLN244 B | 1 |
| ASP153 A | 0.98 |
| CYS160 A | 0.98 |
| ASP153 B | 0.98 |
| VAL148 A | 0.97 |
| GLY149 A | 0.97 |
| ASN151 A | 0.97 |
| GLN244 A | 0.97 |
| ASN277 A | 0.97 |
| VAL148 B | 0.97 |
| GLY149 B | 0.97 |
| ASN151 B | 0.97 |
| CYS160 B | 0.97 |
| GLY29 A | 0.95 |
| VAL35 A | 0.95 |
| CYS38 A | 0.95 |
| ASP92 A | 0.95 |
| THR196 A | 0.95 |
| GLY29 B | 0.95 |
| VAL35 B | 0.95 |
| CYS38 B | 0.95 |
| ASP92 B | 0.95 |
| THR196 B | 0.95 |
| LYS236 B | 0.95 |
| ASN277 B | 0.95 |

**Table S4: Details of Site 1 residues.** Shown are quantile scores in both monomers and the solvent accessible surface area (SASA) to provide an idea of targetability.

| Residue | Quantile Score Chain A | Quantile Score Chain B | SASA [$Å^2$] |
|---|---|---|---|
| Leu30 | 0.98 | 0.98 | 0 |
| Leu32 | 1.00 | 1.00 | 0 |
| Asp33 | 0.95 | 0.97 | 137.86 |
| Asn95 | 0.92 | 0.92 | 13.79 |
| Thr98 | 0.92 | 0.92 | 72.01 |
| Lys100 | 1.00 | 1.00 | 293.05 |
| Tyr101 | 1.00 | 1.00 | 118.61 |
| Phe103 | 0.98 | 0.98 | 128.62 |
| Phe159 | 1.00 | 1.00 | 0 |

**Table S5: Details of Site 2 residues.** Shown are quantile scores in both monomers and the solvent accessible surface area (SASA) to provide an idea of targetability.

| Residue | Quantile Score Chain A | Quantile Score Chain B | SASA [$Å^2$] |
|---|---|---|---|
| Arg4 | 0.97 (B) | 0.97 (A) | 68.49 |
| Arg131 | 1 | 0.98 | 36.50 |
| Asp197 | 0.9 | 0.88 | 101.57 |
| Thr199 | 0.93 | 0.93 | 65.26 |
| Asp289 | 0.98 | 0.98 | 27.98 |
| Glu290 | 0.97 | 0.97 | 25.86 |

**Table S6: Details of Site 3 residues.** Shown are quantile scores in both monomers and the solvent accessible surface area (SASA) to provide an idea of targetability. Highlighted in blue is a cysteine residue which might provide an anchor for drug design.

| Residue | Quantile Score Chain A | Quantile Score Chain B | SASA [Å$^2$] |
|---|---|---|---|
| Lys100 | 0.89 | 0.89 | 145.96 |
| Lys102 | 0.75 | 0.74 | 113.04 |
| Asn151 | 0.97 | 0.97 | 31.34 |
| Ile152 | 0.93 | 0.93 | 5.95 |
| Asp153 | 0.98 | 0.98 | 113.04 |
| Tyr154 | 0.59 | 0.59 | 132.10 |
| Asp155 | 0.92 | 0.92 | 25.18 |
| Cys156 | 1 | 1 | 24.76 |
| Val157 | 0.75 | 0.77 | 0 |
| Ser158 | 0.89 | 0.89 | 15.97 |

**Table S7: Details of Site 4 residues.** Shown are quantile scores in both monomers and the solvent accessible surface area (SASA) to provide an idea of targetability.

| Residue | Quantile Score Chain A | Quantile Score Chain B | SASA [Å$^2$] |
|---|---|---|---|
| Asp33 | 0.93 | 0.93 | 68.93 |
| Asp34 | 0.93 | 0.93 | 45.79 |
| Val35 | 0.95 | 0.95 | 19.56 |
| Tyr37 | 0.85 | 0.85 | 21.65 |
| Arg76 | 0.84 | 0.82 | 170.23 |
| Ile78 | 0.85 | 0.85 | 93.58 |
| Lys90 | 0.82 | 0.82 | 89.29 |
| Val91 | 0.64 | 0.64 | 0.71 |
| Asp92 | 0.95 | 0.95 | 80.04 |
| Thr93 | 0.87 | 0.87 | 60.17 |
| Ala94 | 0.90 | 0.90 | 63.23 |

**Table S8: Scoring of the active site in relation to the 4 identified sites.** The entries are coloured as indicated in the plots in the main results.

| Run | Active Site Score Propensities | Random Site Score [95% CI] | Active Site Score Markov Transients | Random Site Score [95% CI] |
|---|---|---|---|---|
| Site 1 | 0.64 | 0.47 [0.46, 0.47] | 0.69 | 0.53 [0.52, 0.53] |
| Site 2 | 0.49 | 0.48 [0.47, 0.49] | 0.55 | 0.52 [0.52, 0.53] |
| Site 3 | 0.74 | 0.50 [0.49, 0.51] | 0.66 | 0.53 [0.52, 0.53] |
| Site 4 | 0.43 | 0.46 [0.46, 0.47] | 0.52 | 0.50 [0.50, 0.51] |

**Table S9: Scoring of the active site in relation to the small fragments in close proximity of the identified sites.** The entries are coloured according to which site the respective fragment is close to.

| Run | Active Site Score Propensities | Random Site Score [95% CI] | Active Site Score Markov Transients | Random Site Score [95% CI] |
|---|---|---|---|---|
| 5RED 1 | 0.63 | 0.49 [0.48, 0.49] | 0.65 | 0.54 [0.54, 0.55] |
| 5REI 1 | 0.73 | 0.48 [0.47, 0.49] | 0.57 | 0.56 [0.55, 0.56] |
| 5RGJ 1 | 0.65 | 0.46 [0.46, 0.47] | 0.65 | 0.53 [0.53, 0.54] |
| 5RGR 1 | 0.67 | 0.49 [0.48, 0.50] | 0.67 | 0.54 [0.54, 0.55] |
| 5RF5 1 | 0.65 | 0.48 [0.47, 0.49] | 0.62 | 0.54 [0.53, 0.54] |
| 5RF0 2 | 0.44 | 0.52 [0.51, 0.53] | 0.56 | 0.55 [0.54, 0.56] |
| 5RGQ 2 | 0.35 | 0.49 [0.48, 0.50] | 0.52 | 0.52 [0.51, 0.52] |
| 5RE5 4 | 0.40 | 0.49 [0.48, 0.49] | 0.51 | 0.53 [0.53, 0.54] |
| 5RE7 4 | 0.53 | 0.45 [0.44, 0.46] | 0.58 | 0.50 [0.49, 0.51] |
| 5RFC 4 | 0.42 | 0.48 [0.47, 0.48] | 0.49 | 0.52 [0.52, 0.53] |
| 5RGG 4 | 0.24 | 0.51 [0.50, 0.52] | 0.41 | 0.53 [0.53, 0.54] |
| 5RF9 1 3 | 0.69 | 0.50 [0.49, 0.51] | 0.69 | 0.54 [0.53, 0.54] |
| 5RE8 1 4 | 0.70 | 0.47 [0.46, 0.48] | 0.74 | 0.50 [0.49, 0.51] |
| 5RF4 1 4 | 0.64 | 0.45 [0.45, 0.46] | 0.67 | 0.51 [0.50, 0.51] |
| 5RFD 1 4 | 0.64 | 0.47 [0.46, 0.47] | 0.58 | 0.52 [0.52, 0.53] |
